# TrIPP: a Trajectory Iterative p*K*_*a*_ Predictor

**DOI:** 10.1101/2025.09.02.673559

**Authors:** Christos Matsingos, Ka Fu Man, Arianna Fornili

## Abstract

The protonation propensity of ionisable residues in proteins can change in response to changes in the local residue environment. The link between protein dynamics and p*K*_a_ is particularly important in pH regulation of protein structure and function. Here, we introduce TrIPP (Trajectory Iterative p*K*_a_ Predictor), a Python tool to monitor and analyse changes in the p*K*_a_ of ionisable residues during Molecular Dynamics simulations of proteins. We show how TrIPP can be used to identify residues with physiologically relevant variations in their predicted p*K*_a_ values during the simulations, and link them to changes in the local and global environment. TrIPP is available at https://github.com/fornililab/TrIPP.

## 1 Introduction

The conformation of proteins can depend on different factors, including the pH. Indeed, several examples exist of proteins that undergo structural shifts in response to physiological pH changes (Schönichen *et al*., 2013; Di Russo *et al*., 2012; Matsingos *et al*., 2024; Warwicker, 2022). The link between pH and protein structure is mediated by the dependence between p*K*_a_ values, which regulate the protonation propensity of ionisable residues, and the local residue environment, which can significantly change with the protein conformation.

Due to the difficulty in experimentally measuring p*K*_a_ values, different computational methods (Alexov *et al*., 2011; Olsson *et al*., 2011; Wei *et al*., 2023; Gordon *et al*., 2005; Anandakrishnan *et al*., 2012; Rabenstein and Knapp, 2001; Reis *et al*., 2020) have been developed to predict p*K*_a_ deviations from their reference values (isolated amino acid in solution). However, these approaches are normally applied to single structures, for example to define the protonation state of ionisable residues before starting a Molecular Dynamics (MD) simulation. Here, we extend this framework by introducing TrIPP (Trajectory Iterative p*K*_a_ Predictor), a tool to monitor and analyse p*K*_a_ changes during MD trajectories.

TrIPP is based on the iterative use of PROPKA 3 (Olsson *et al*., 2011), one of the most popular approaches for single-structure p*K*_a_ prediction. We show that TrIPP can be used to visualise p*K*_a_ distributions and time evolutions over MD trajectories and identify residues with conformation-dependent p*K*_a_ values. Pseudo-mutations can be carried out to assess the influence of specific residues on p*K*_a_ regulation. In addition, TrIPP can cluster trajectory frames according to the local environment of selected ionisable residues, to identify structures representative of the environments leading to different p*K*_a_ values. Insights from TrIPP can be used to guide further studies and formulate mechanistic hypotheses on pH-dependent regulation of protein function.

## 2 Implementation

TrIPP has been implemented using Python 3.9 along with several packages, including PROPKA 3.5.0 (Olsson *et al*., 2011), MDAnalysis 2.5.0 (Michaud-Agrawal *et al*., 2011; Gowers *et al*., 2016), Pandas 2.0.3 (McKinney, 2010), NumPy 1.25.0 (Harris *et al*., 2020), scikit-learn 1.5.0 (Pedregosa *et al*., 2011), scikit-learn-extra 0.3.0, scipy 1.13.0 (Virtanen *et al*., 2020), and seaborn 0.13.2 (Waskom, 2021). The software has been packaged with Poetry 1.7.1 and is available for installation through PyPI.

### 2.1 TrIPP workflow

The workflow comprises four stages: data input, data pre-processing, p*K*_a_ prediction, and data analysis (Fig. 1A).

**Figure 1.**
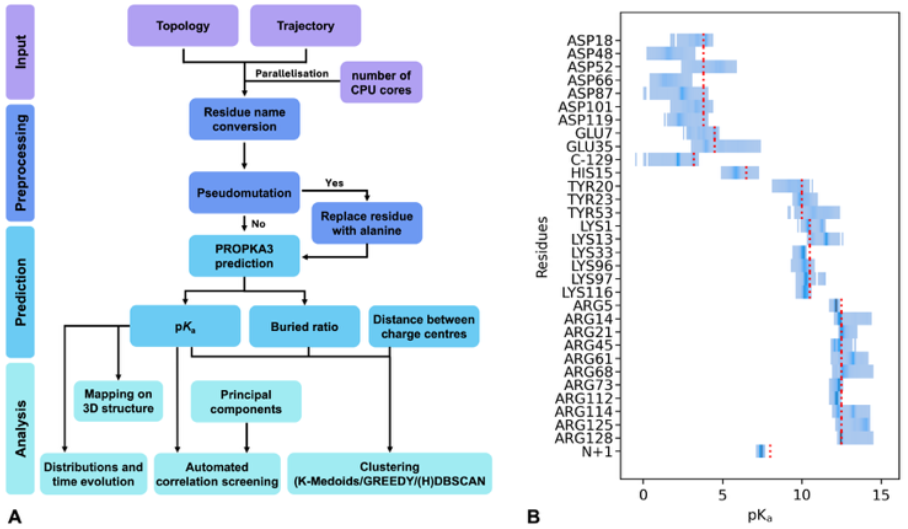
TrIPP overview and application to the hen egg white lysozyme. **(A)** Different stages in the TrIPP workflow. **(B)** Distributions of predicted p*K*_a_ values for ionisable residues in lysozyme during one of the trajectories (MD1). For each residue, p*K*_a_ values are binned along the x-axis and more frequent values are indicated with a darker shading. A red vertical line indicates the PROPKA model value for the corresponding residue type.

#### Data Input

TrIPP uses MDAnalysis to load input trajectory and topology files. Parallelisation is carried out by splitting each trajectory into as many subsets of frames as the number of cores used in the calculation. Each subset is processed by a different core in the subsequent steps.

#### Data Preprocessing

Preprocessing includes an optional pseudo-mutation feature, which can be activated by the user to replace selected residues with alanine. Where required, residue names are also converted into versions recognised by PROPKA. Each snapshot is then temporarily converted to a PDB file, which is the format required by PROPKA.

#### Prediction

PROPKA is iteratively run on each snapshot and its output is parsed to extract the p*K*_a_ values and (optionally) the buried ratio (given as %) of all the ionisable residues. Temporary files (PDB snapshot and PROPKA output) are deleted after processing. The extracted values are stored in CSV files and sorted by snapshot timeframe.

#### Data analysis

Plots of p*K*_a_ distributions and time evolutions can be generated from the p*K*_a_ values predicted for each snapshot. TrIPP can write PyMOL sessions (PSE format) to colour-map different properties on the 3D structure, including the p*K*_a_ values of all the ionisable residues for a selected frame or averaged over a set of trajectories. The difference between the average and PROPKA model p*K*_a_ values can be also mapped. If the projection of the trajectories on selected principal components is provided by the user, an automatic scanning of the correlation coefficients between projections and p*K*_a_ values can be run to identify possible p*K*_a_-collective motions coupling.

The generated data can be used to extract representative structures from the trajectory using one of the available clustering algorithms: K-Medoids^10^, greedy (Micheletti *et al*., 2000), DBSCAN (Ester *et al*., 1996), and HDBSCAN (Campello *et al*., 2013). The feature matrix used for the clustering is built from p*K*_a_ values of user-selected residues and (optionally) their buried ratio and inter-residue distances. Features are Z-score normalised before clustering and (optionally) subjected to Principal Component Analysis (PCA). For each method, clustering hyperparameters can be optionally optimised via grid search, using silhouette widths (Rousseeuw, 1987) to assess the quality of the clustering.

At last, TrIPP provides a function to automatically scan all ionisable residues for possible correlations between their p*K*_a_ values and a time-dependent property provided by the user. For example, projections from a principal component analysis of the trajectories can be used to represent collective motions in the protein.

### 2.2 TrIPP classes

TrIPP provides its functionalities through 3 main classes: *Trajectory* (input, preprocessing and prediction), *Visualisation* (3D mapping of p*K*_a_-related values) and *Clustering*. A comprehensive tutorial demonstrating the use of these classes is included in the TrIPP github distribution as Jupyter notebook.

## 3 Application

TrIPP was applied to MD simulations of the hen egg-white lysozyme to illustrate its features. This enzyme was chosen since the p*K*_a_ values of its residues have been extensively studied both experimentally and computationally (Williams *et al*., 2010; Webb *et al*., 2011). In particular, the p*K*_a_ of acidic residues in its active site has been shown to be modulated by interactions with nearby residues (Williams *et al*., 2010). MD simulations were run for 300 ns (production) in five replicates (MD1 to MD5), system preparation and MD protocol are described in the Supplementary Methods and Supplementary Table S1. TrIPP was run on each replica using frames sampled every 100 ps.

Average p*K*_a_ values show small variations across replicas, indicating good reproducibility, and a good agreement with the available experimental values (Supplementary Table S2). Inspecting the distribution of p*K*_a_ values observed for all ionizable residues (Fig. 1B and Supplementary Figure S1) highlights the residues that have p*K*_a_ values (blue shades) significantly shifted from their model p*K*_a_ (dotted red line). Among the acidic residues, Asp52 and Glu35 stand out because their p*K*_a_ values are upshifted towards the region of physiological pH. This can be also easily spotted in the PyMOL visualisations of the average p*K*_a_ values for acidic residues and their deviation from model values (Supplementary Figure S2), where Asp52 and Glu35 are the only acidic residues with an increase in the average p*K*_a_ from the model values. While the average increase is small, inspection of the p*K*_a_ time evolution for these residues indicates that values can be as high as ∼ 7 for Glu35 and almost 6 for Asp52 (Fig. 2A). Interestingly, Glu35 and Asp52 are located in the lysozyme active site and are directly involved in the catalytic mechanism, where Glu35 changes its protonation state (Ramos *et al*., 2025).

**Figure 2.**
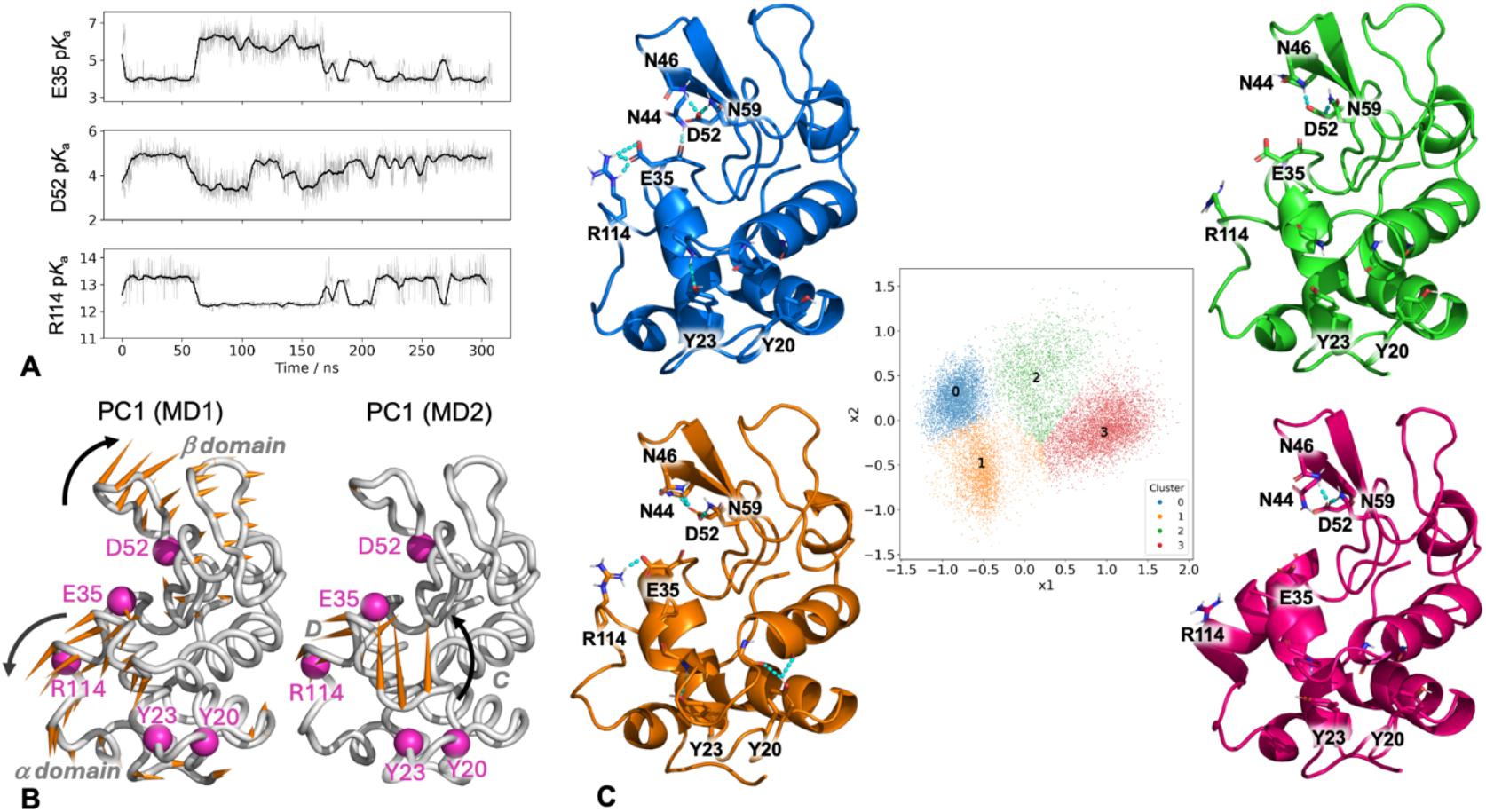
TrIPP-based analysis of MD simulations of lysozyme. **(A)** Time evolution of p*K*_a_ values predicted for Glu35, Asp52 and Arg114 during one of the lysozyme trajectories (MD1). The running average (5-ns window, dark shade) is shown together with the instantaneous values (light shade). **(B)** Porcupine representation of the first Principal Component (PC1) from the MD1 and MD2 lysozyme trajectories. Orange spikes show the direction and relative amplitude of motion of each residue along the PC (spikes shorter than 20% of the maximum length are omitted). Key residues are labelled and their position indicated with magenta spheres. Additional labels indicate the α and β domains the left panel, and the C and D helices in the right panel. **(C)** K-Medoid cluster representatives (cartoon) illustrating different structural environments leading to different p*K*_a_ values for Glu35, Asp52, Arg114, Tyr20, and Tyr23. Key residues are highlighted as sticks and labelled. Polar contacts from PyMOL are represented with dashed lines. The feature matrix used for clustering was first subjected to dimensionality reduction via a Principal Component Analysis. The middle panel shows a projection of all trajectories on the first two principal components x_1_ and x_2_, with each frame coloured according to its cluster ID.

To detect possible couplings between p*K*_a_ changes and protein dynamics, the correlation coefficient between the p*K*_a_ time evolution and the main collective motions observed during the simulations (as described by projections of each trajectory on its top-ranking principal components) was calculated for each ionizable residue (Supplementary Figure S3). The largest correlations (0.5 or higher) with the first two principal components (PC1 and PC2) were observed for Tyr20, Glu35, Asp52, Arg114, and to a less extent Tyr23, suggesting that the dominant protein motions can modulate the p*K*_a_ value of these residues.

Two different types of recurring motions were identified. The first one, exemplified by PC1 from MD1 (Fig. 2B, left), is the previously observed hinge-bending motion of the α- and β-domains(Brooks and Karplus, 1985), where Asp52, Glu35, Arg114 are part of or close to the moving regions. The projections of the different trajectories (Supplementary Figure S4A) show patterns that are clearly correlated (Glu35) or anti-correlated (Asp52, Arg114) with p*K*_a_ time evolutions (Supplementary Figure S5). PC1 from MD2 instead describes a more localised restructuring of the loop between helices C and D, which is positioned above Tyr20 and Tyr23 (Fig. 2B, right).

Trajectories were then clustered to identify structures representative of the different environments around residues with motion-sensitive p*K*_a_ values (Fig. 2C). Clusters 0 and 1 feature a salt-bridge between Glu35 and Arg114, which is instead absent in clusters 2 and 3. This explains their distinct p*K*_a_ distributions across the clusters (Supplementary Figure S6), since acidic and basic residues forming salt bridges are expected to have down- and up-shifted p*K*_a_ values, respectively, compared to their non-interacting states. Clusters 1 differs from cluster 0 because of the presence of additional hydrogen bonding between Asp52 and Asn44, which explains the lower Asp52 p*K*_a_ values for this cluster, and between Tyr20 and the CD loop backbone, which leads to increased Tyr20 p*K*_a_ values.

The environment of Glu35 and Asp52 was further analysed using the pseudo-mutation tool implemented in TrIPP, where p*K*_a_ values are re-calculated after replacing selected residues with alanine, while keeping all the other coordinates in the protein unchanged. Replacing residues Asn44, Asn46 and Asn59 with alanine shows an increase of Asp52 p*K*_a_ across all the replicas (Supplementary Figure S7A and C). This is consistent with the formation of hydrogen bonds between all these residues and Asp52. Interestingly, instead of just shifting the Asp52 p*K*_a_ time evolution profile, Asn44 also modulates its shape. Indeed, the Asn44Ala pseudo-mutant shows smaller p*K*_a_ fluctuations compared to the wild type (Figure Supplementary Figure S7C), consistently with the formation and breaking of Asn44-Asp52 hydrogen bonds during the trajectory. The role of Asn44 in stabilising Asp52 negative charge and modulating its availability for substrate binding has been highlighted in recent structural studies of lysozyme (Ramos *et al*., 2025). Similarly, replacing Arg114 with alanine induces an increase of the Asp52 p*K*_a_ but only in the parts of the trajectory where the two residues are interacting with each other (Supplementary Figure S7B and D). These examples illustrate how analysing the effect of pseudo-mutations on the shape of the p*K*_a_ time evolution profile of a given residue can help identifying the residues that mediate the dynamic response of its p*K*_a_ value.

## 4 Concluding remarks

We developed a Python tool to monitor, analyse and visualise changes in p*K*_a_ values during MD simulations. Information from TrIPP can be used to guide protein modelling, especially when a pH sensing behaviour is known or suspected (Matsingos *et al*., 2024). It is important to highlight that TrIPP is not meant to replace in any way constant-pH simulations (Williams *et al*., 2010; Oliveira *et al*., 2022), where the protonation state of ionisable residues is allowed to change. Here, we have shown how TrIPP can be used in conjunction with conventional MD simulations. While the sampled frames will reflect fixed protonation states, detection of p*K*_a_ changes during the trajectory can highlight residues that might require further simulations with alternative protonation states. Observing large correlations between residues p*K*_a_ and global motions might suggest that pH modulation of protein conformation is involved, warranting further investigation. Moreover, clustering and pseudo-mutation analyses can be used to build testable hypotheses about the molecular mechanisms underlying pH-dependent structural and functional changes.

## Acknowledgements

We would like to thank Yu-Yuan Yang and Dr Alessandro Pandini for testing and providing feedback on earlier versions of TrIPP.

## References

Alexov, E., Mehler, E.L., Baker, N., et al. (2011) Progress in the prediction of p Ka values in proteins. Proteins, 79, 3260–3275.

Anandakrishnan, R., Aguilar, B., and Onufriev, A.V. (2012) H++ 3.0: automating pK prediction and the preparation of biomolecular structures for atomistic molecular modeling and simulations. Nucleic Acids Research, 40, W537–W541.

Brooks, B. and Karplus, M. (1985) Normal modes for specific motions of macro-molecules: application to the hinge-bending mode of lysozyme. Proc. Natl. Acad. Sci. U.S.A., 82, 4995–4999.

Campello, R.J.G.B., Moulavi, D., and Sander, J. (2013) Density-Based Clustering Based on Hierarchical Density Estimates. In, Pei, J., Tseng, V.S., Cao, L., et al. (eds), Advances in Knowledge Discovery and Data Mining. Springer Berlin Heidelberg, Berlin, Heidelberg, pp. 160–172.

Di Russo, N.V., Estrin, D.A., Martí, M.A., et al. (2012) pH-Dependent Conformational Changes in Proteins and Their Effect on Experimental pKas: The Case of Nitrophorin 4. PLoS Comput Biol, 8, e1002761.

Ester, M., Kriegel, H.-P., Sander, J., et al. (1996) A density-based algorithm for discovering clusters in large spatial databases with noise. In, Proceedings of the Second International Conference on Knowledge Discovery and Data Mining, KDD’96. AAAI Press, Portland, Oregon, pp. 226–231.

Gordon, J.C., Myers, J.B., Folta, T., et al. (2005) H++: a server for estimating p Ka s and adding missing hydrogens to macromolecules. Nucleic Acids Research, 33, W368–W371.

Gowers, R., Linke, M., Barnoud, J., et al. (2016) MDAnalysis: A Python Package for the Rapid Analysis of Molecular Dynamics Simulations. Austin, Texas, pp. 98– 105.

Harris, C.R., Millman, K.J., Van Der Walt, S.J., et al. (2020) Array programming with NumPy. Nature, 585, 357–362.

Matsingos, C., Howell, L.A., McCormick, P.J., et al. (2024) Elucidating the Activation Mechanism of the Proton-sensing GPR68 Receptor. Journal of Molecular Biology, 436, 168688.

McKinney, W. (2010) Data Structures for Statistical Computing in Python. Austin, Texas, pp. 56–61.

Michaud-Agrawal, N., Denning, E.J., Woolf, T.B., et al. (2011) MDAnalysis: A toolkit for the analysis of molecular dynamics simulations. J Comput Chem, 32, 2319–2327.

Micheletti, C., Seno, F., and Maritan, A. (2000) Recurrent oligomers in proteins: An optimal scheme reconciling accurate and concise backbone representations in automated folding and design studies. Proteins: Structure, Function, and Bioinformatics, 40, 662–674.

Oliveira, V.M. de, Liu, R., and Shen, J. (2022) Constant pH molecular dynamics simulations: Current status and recent applications. Current Opinion in Structural Biology, 77, 102498.

Olsson, M.H.M., Søndergaard, C.R., Rostkowski, M., et al. (2011) PROPKA3: Consistent Treatment of Internal and Surface Residues in Empirical p K a Predictions. J. Chem. Theory Comput., 7, 525–537.

Pedregosa, F., Varoquaux, G., Gramfort, A., et al. (2011) Scikit-learn: Machine Learning in Python. Journal of Machine Learning Research, 12, 2825–2830.

Rabenstein, B. and Knapp, E.-W. (2001) Calculated pH-Dependent Population and Protonation of Carbon-Monoxy-Myoglobin Conformers. Biophysical Journal, 80, 1141–1150.

Ramos, J., Laux, V., Mason, S.A., et al. (2025) Structure and dynamics of the active site of hen egg-white lysozyme from atomic resolution neutron crystallography. Structure, 33, 136-148.e3.

Reis, P.B.P.S., Vila-Viçosa, D., Rocchia, W., et al. (2020) PypKa: A Flexible Python Module for Poisson–Boltzmann-Based pKa Calculations. J. Chem. Inf. Model., 60, 4442–4448.

Rousseeuw, P.J. (1987) Silhouettes: A graphical aid to the interpretation and validation of cluster analysis. Journal of Computational and Applied Mathematics, 20, 53–65.

Schönichen, A., Webb, B.A., Jacobson, M.P., et al. (2013) Considering Protonation as a Posttranslational Modification Regulating Protein Structure and Function. Annu. Rev. Biophys., 42, 289–314.

Virtanen, P., Gommers, R., Oliphant, T.E., et al. (2020) SciPy 1.0: fundamental algorithms for scientific computing in Python. Nat Methods, 17, 261–272.

Warwicker, J. (2022) The Physical Basis for pH Sensitivity in Biomolecular Structure and Function, With Application to the Spike Protein of SARS-CoV-2. Front. Mol. Biosci., 9, 834011.

Waskom, M. (2021) seaborn: statistical data visualization. JOSS, 6, 3021.

Webb, H., Tynan-Connolly, B.M., Lee, G.M., et al. (2011) Remeasuring HEWL pKa values by NMR spectroscopy: Methods, analysis, accuracy, and implications for theoretical pKa calculations. Proteins, 79, 685–702.

Wei, W., Hogues, H., and Sulea, T. (2023) Comparative Performance of High-Throughput Methods for Protein p Ka Predictions. J. Chem. Inf. Model., 63, 5169–5181.

Williams, S.L., De Oliveira, C.A.F., and McCammon, J.A. (2010) Coupling Constant pH Molecular Dynamics with Accelerated Molecular Dynamics. J. Chem. Theory Comput., 6, 560–568.

