## Supplementary Material for "TrIPP: a Trajectory Iterative p*K*_*a*_ Predictor"

### Supplementary Methods

#### Molecular Dynamics Simulations

Molecular Dynamics (MD) simulations of the hen egg-white lysozyme were performed with GROMACS[1] (version 2020.3[2]) using the Amber99SB*-ILDN[3] force field. The starting structure was prepared from the PDB crystal structure 1AKI[4]. Water molecules were removed and the protonation state of ionisable residues at pH 7.0 was predicted with PROPKA 3 (version 3.4.0)[5]. The protein was solvated with a truncated octahedral box with a 12-Å minimum distance between the walls of the box and the protein, resulting in about 9,500 TIP3P[6] water molecules. Counterions were added to neutralise the system and additional $Na^+$ and $Cl^-$ ions were introduced to reach an overall ionic strength of 0.05 $molL^{-1}$. The total number of atoms was 30,425.

Periodic boundary conditions were applied in all three directions. The Particle Mesh Ewald (PME) method was used for electrostatic interactions, with a 9-Å cutoff for the direct space sums, a 1.2-Å FFT grid spacing, and a 4-th order interpolation polynomial for the reciprocal space sums. A 9-Å cutoff was used for van der Waals interactions and long-range dispersion corrections were applied.

The system was energy-minimised in 3 stages: 1. up to 5,000 steps of steepest descent with positional restraints on heavy atoms (4.8-kcal/mol/$Å^2$ force constant), 2. up to 5,000 steps of steepest descent with no restraints, and 3. up to 2,000 steps of conjugate gradient (no restraints).

Five sets of replicas were started from the energy-minimised system. Equilibration was run in different stages as indicated in Table S1. Positional restraints on heavy atoms were initially set to 4.8 kcal/mol/$Å^2$ and gradually decreased to 0 (Steps 1 to 6 in Table S1). The time step was set to 1 fs in the first 9 equilibration stages and increased to 2 fs in the last one. The production phase was run for 300 ns using the same conditions as the last equilibration step. The LINCS[7] algorithm was used to constrain all covalent bonds to hydrogen atoms in the protein, while SETTLE[8] was used for water molecules. Frames were saved every 100 ps.

#### Analysis of trajectories

Trajectories were analysed using TrIPP as described in the main text. Additional details are reported in the following for cluster analysis and collective motion determination.

Cluster analysis was performed using the TrIPP Clustering class. The feature matrix was built using Z-score normalised p$K_a$ values and distances between the charge centres of Tyr20, Tyr23, Glu35, Asp52, Arg114 for all the frames from the five replicas. Dimensionality reduction via Principal Component Analysis (PCA) was applied and the top components explaining at least 90% of the cumulative variance were retained. Clusters were determined with the K-Medoids method, using euclidean distances and setting the number of clusters to four. The medoids were extracted as the representative structures.

Collective motions were determined and analysed using the GROMACS tools *gmx covar* and *gmx anaeig*. A PCA was performed on the C$\alpha$ atom coordinates of each trajectory (production part only). The trajectories were then projected onto their top 3 Principal Components (PCs) and TrIPP was used to automatically scan all ionisable residues for possible correlations (Pearson correlation coefficient) between their p$K_a$ values and each projection. PCs were also compared to identify recurring motions across the replicas. Porcupine plots were generated to visualise the type of motions represented by each PC and inner products between all possible pairs of PCs were calculated.

**Table S1.** Full description of the equilibration stages.

| Step | Duration (ns) | Conditions | Temperature (K) | Restraints / kcal/mol/Å$^2$ |
|---|---|---|---|---|
| 1 | 0.1 | NVT[a] | 200 | 4.8[d] |
| 2 | 0.1 | NVT[a] | 250 | 2.4[d] |
| 3 | 0.1 | NVT[a] | 300 | 2.4[d] |
| 4 | 0.1 | NVT[a] | 300 | 1.2[d] |
| 5 | 0.1 | NVT[a] | 300 | 0.6[d] |
| 6 | 1 | NVT[a] | 300 | 0.6[e] |
| 7 | 1 | NVT[a] | 300 | - |
| 8 | 2 | NPT[b] | 300 | - |
| 9 | 2 | NPT[c] | 300 | - |
| 10 | 2 | NPT[c, f] | 300 | - |

[a]Berendsen thermostat[9] (0.2-ps coupling constant)

[b]Berendsen thermostat (0.2-ps coupling constant) and Berendsen barostat (1-ps coupling constant, p = 1 bar).

[c]V-rescale thermostat[10] (0.1-ps coupling constant) and Parrinello-Rahman barostat[11] (2-ps coupling constant, p = 1 bar).

[d]Restraint on all protein heavy atoms.

[e]Restraint on protein backbone heavy atoms.

[f]2-fs time step

**Table S2.** Experimental and average predicted p$K_a$ values for ionisable residues in lysozyme.

| Residue | Bartik et al[12] | Webb et al.[13] | PROPKA3[a] | MD1[b] | MD2[b] | MD3[b] | MD4[b] | MD5[b] |
|---|---|---|---|---|---|---|---|---|
| **N-term** | - | - | 7.4 | 7.4 ± 0.1 | 7.4 ± 0.1 | 7.4 ± 0.1 | 7.5 ± 0.1 | 7.4 ± 0.1 |
| **LYS1** | - | - | 11.2 | 10.7 ± 0.4 | 10.8 ± 0.4 | 10.7 ± 0.4 | 10.7 ± 0.4 | 10.7 ± 0.4 |
| **ARG5** | - | - | 12.1 | 12.1 ± 0.1 | 12.2 ± 0.1 | 12.2 ± 0.1 | 12.2 ± 0.1 | 12.2 ± 0.1 |
| **GLU7** | 2.85 ± 0.25 | 2.6 ± 0.2 | 2.9 | 3.8 ± 0.6 | 3.7 ± 0.6 | 3.7 ± 0.6 | 3.8 ± 0.6 | 3.8 ± 0.6 |
| **LYS13** | - | - | 10.6 | 11.3 ± 0.4 | 11.3 ± 0.4 | 11.2 ± 0.4 | 11.3 ± 0.4 | 11.3 ± 0.4 |
| **ARG14** | - | - | 12.3 | 12.5 ± 0.4 | 12.6 ± 0.5 | 12.5 ± 0.4 | 12.6 ± 0.5 | 12.4 ± 0.2 |
| **HIS15** | 5.36 ± 0.07 | 5.5 ± 0.2 | 6.3 | 5.8 ± 0.3 | 5.9 ± 0.3 | 5.9 ± 0.3 | 5.9 ± 0.3 | 5.8 ± 0.2 |
| **ASP18** | 2.66 ± 0.08 | 2.8 ± 0.3 | 3.3 | 3.5 ± 0.4 | 3.5 ± 0.4 | 3.5 ± 0.5 | 3.5 ± 0.4 | 3.5 ± 0.4 |
| **TYR20** | - | - | 9.6 | 9.5 ± 0.5 | 10.0 ± 0.5 | 9.8 ± 0.7 | 9.6 ± 0.6 | 9.4 ± 0.5 |
| **ARG21** | - | - | 12.8 | 12.6 ± 0.2 | 12.8 ± 0.3 | 12.7 ± 0.3 | 13.1 ± 0.5 | 12.7 ± 0.3 |
| **TYR23** | - | - | 9.9 | 10.1 ± 0.3 | 10.4 ± 0.3 | 10.2 ± 0.3 | 10.2 ± 0.3 | 10.0 ± 0.3 |
| **LYS33** | - | - | 10.2 | 10.1 ± 0.1 | 10.1 ± 0.1 | 10.0 ± 0.1 | 10.1 ± 0.1 | 10.1 ± 0.1 |
| **GLU35** | 6.20 ± 0.10 | 6.1 ± 0.4 | 5.8 | 4.8 ± 1.0 | 5.0 ± 0.7 | 4.5 ± 0.7 | 4.9 ± 0.7 | 4.9 ± 1.0 |
| **ARG45** | - | - | 12.0 | 12.3 ± 0.2 | 12.2 ± 0.2 | 12.3 ± 0.2 | 12.3 ± 0.2 | 12.2 ± 0.2 |
| **ASP48** | <2.5 | 1.4 ± 0.2 | 1.6 | 1.7 ± 0.4 | 1.8 ± 0.4 | 1.8 ± 0.4 | 1.9 ± 0.5 | 1.9 ± 0.4 |
| **ASP52** | 3.68 ± 0.08 | 3.6 ± 0.3 | 3.3 | 4.3 ± 0.6 | 3.9 ± 0.5 | 4.2 ± 0.6 | 4.1 ± 0.6 | 3.9 ± 0.6 |
| **TYR53** | - | - | 11.9 | 11.0 ± 0.5 | 10.8 ± 0.5 | 10.9 ± 0.5 | 10.7 ± 0.5 | 11.0 ± 0.5 |
| **ARG61** | - | - | 13.3 | 13.1 ± 0.4 | 13.0 ± 0.4 | 13.0 ± 0.3 | 12.9 ± 0.5 | 12.9 ± 0.4 |
| **ASP66** | <2.0 | 1.2 ± 0.2 | 1.5 | 1.6 ± 0.4 | 1.5 ± 0.4 | 1.6 ± 0.4 | 1.5 ± 0.4 | 1.6 ± 0.4 |
| **ARG68** | - | - | 12.5 | 12.7 ± 0.3 | 12.7 ± 0.3 | 12.8 ± 0.3 | 12.7 ± 0.3 | 12.8 ± 0.3 |
| **ARG73** | - | - | 12.3 | 12.3 ± 0.1 | 12.2 ± 0.1 | 12.2 ± 0.2 | 12.2 ± 0.1 | 12.3 ± 0.3 |
| **ASP87** | 2.07 ± 0.15 | 2.2 ± 0.1 | 2.6 | 2.6 ± 0.4 | 2.4 ± 0.6 | 2.5 ± 0.4 | 2.5 ± 0.5 | 2.7 ± 0.5 |
| **LYS96** | - | - | 10.0 | 10.2 ± 0.2 | 10.1 ± 0.2 | 10.2 ± 0.2 | 10.1 ± 0.2 | 10.2 ± 0.2 |

| | | | | | | | | |
|---|---|---|---|---|---|---|---|---|
| **LYS97** | - | - | 10.3 | 10.3 ± 0.3 | 10.2 ± 0.1 | 10.3 ± 0.3 | 10.5 ± 0.4 | 10.2 ± 0.1 |
| **ASP101** | 4.09 ± 0.07 | 4.5 ± 0.1 | 4.1 | 3.3 ± 0.5 | 3.4 ± 0.3 | 3.3 ± 0.4 | 3.3 ± 0.6 | 3.4 ± 0.5 |
| **ARG112** | - | - | 12.2 | 12.2 ± 0.1 | 12.2 ± 0.1 | 12.2 ± 0.1 | 12.2 ± 0.1 | 12.2 ± 0.1 |
| **ARG114** | - | - | 12.2 | 12.8 ± 0.5 | 12.7 ± 0.6 | 12.9 ± 0.5 | 12.7 ± 0.5 | 12.8 ± 0.6 |
| **LYS116** | - | - | 10.2 | 10.2 ± 0.1 | 10.2 ± 0.1 | 10.2 ± 0.1 | 10.2 ± 0.1 | 10.2 ± 0.1 |
| **ASP119** | 3.20 ± 0.09 | 3.5 ± 0.3 | 3.0 | 2.5 ± 0.5 | 2.4 ± 0.3 | 2.5 ± 0.4 | 2.4 ± 0.4 | 2.4 ± 0.3 |
| **ARG125** | - | - | 13.4 | 13.6 ± 0.6 | 13.8 ± 0.4 | 13.7 ± 0.4 | 13.8 ± 0.4 | 13.8 ± 0.4 |
| **ARG128** | - | - | 12.6 | 12.5 ± 0.2 | 12.5 ± 0.2 | 12.6 ± 0.5 | 12.5 ± 0.2 | 12.5 ± 0.2 |
| **C-term** | 2.75 ± 0.02 | 2.7 ± 0.2/<br>3.9 ± 0.1 | 3.0 | 2.4 ± 0.6 | 2.4 ± 0.6 | 2.7 ± 0.6 | 2.4 ± 0.6 | 2.5 ± 0.5 |

[a]Values are calculated on the initial energy-minimised structure.

[b]Average values calculated over the MD1-5 replicas (production) ± standard deviation.

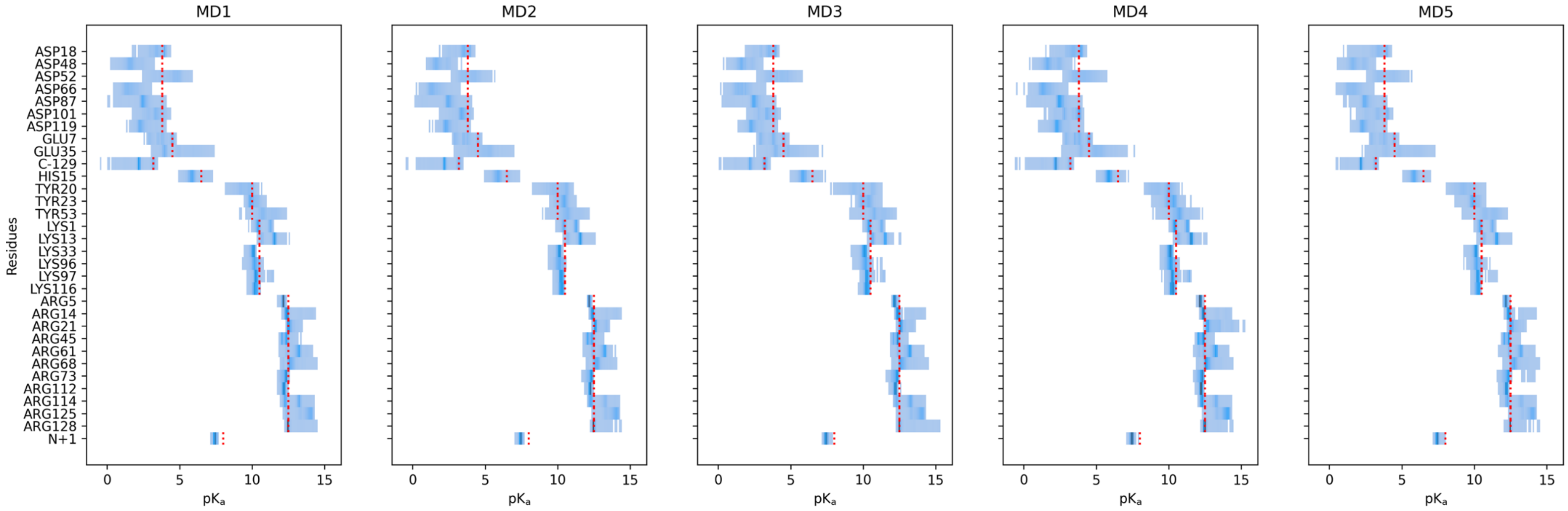


**Figure S1.** Distributions of predicted p$K_a$ values for ionisable residues in lysozyme during the MD1-5 trajectories. For each residue, p$K_a$ values are binned along the x-axis and more frequent values are indicated with a darker shading. A red vertical line indicates the PROPKA model value for the corresponding residue type.

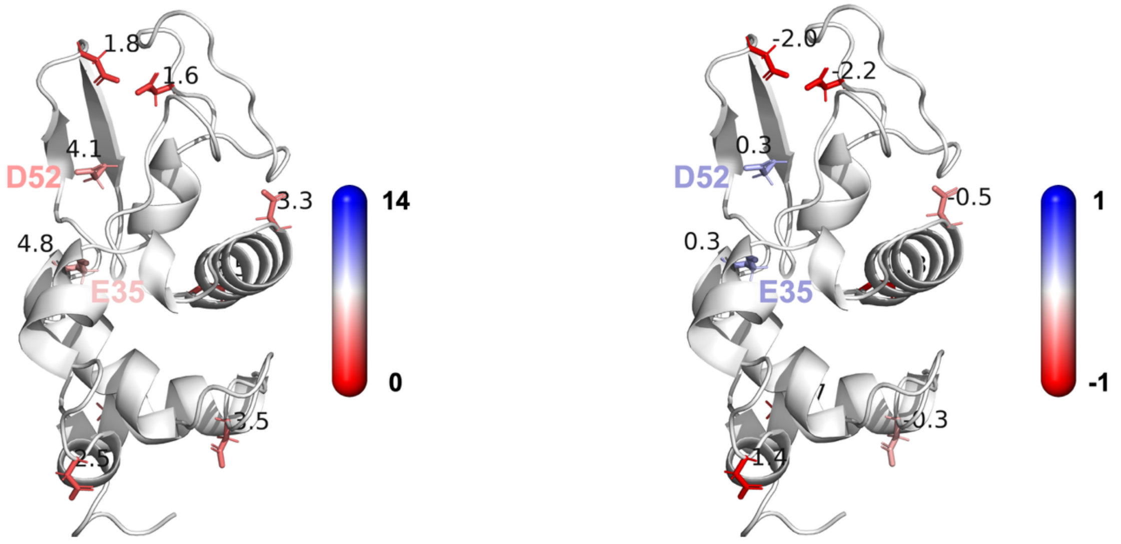


**Figure S2.** Colour mapping of average p$K_a$ values (left) and of the difference between average and PROPKA model values (right) for acidic residues in lysozyme. Only Glu35 and Asp52 are labelled.

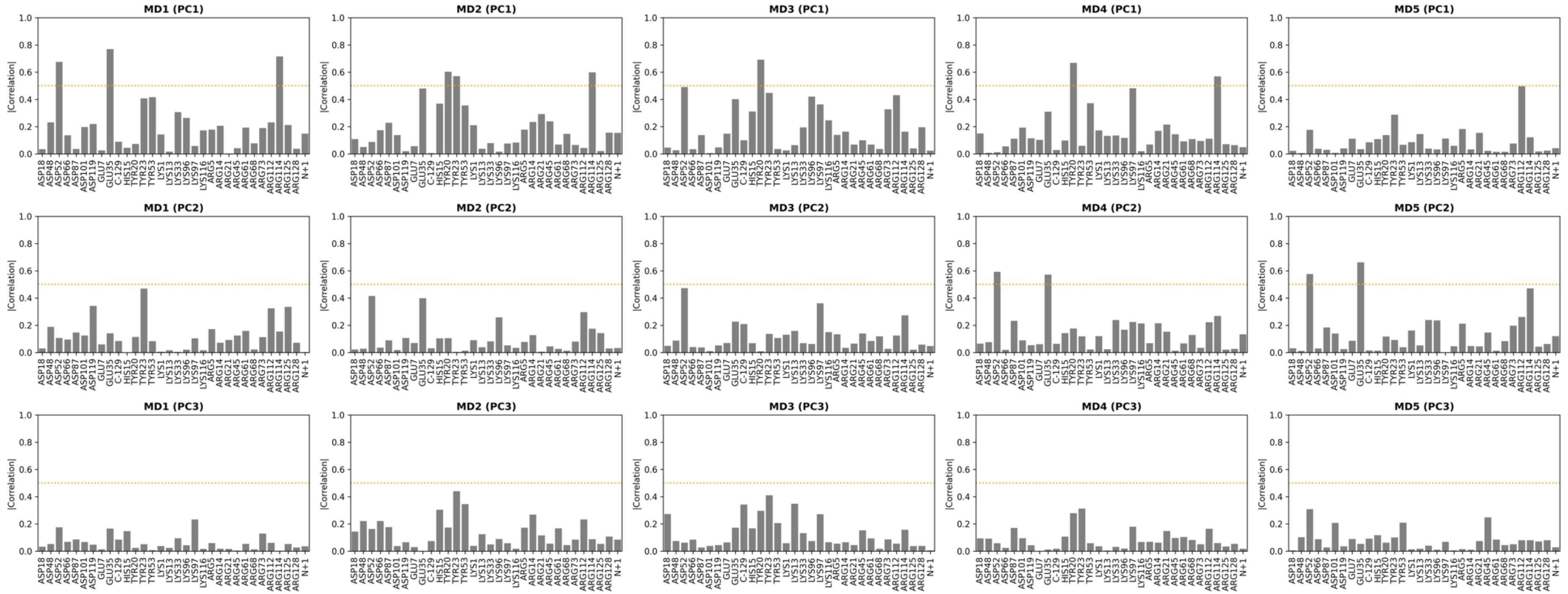


**Figure S3.** Bar plots of Pearson correlation coefficients (absolute value) between the time evolution of $pK_a$ values predicted for all ionisable residues in lysozyme and the projection of each replica (MD1 to MD5) onto its first three principal components (PCs). The N-terminal amino group and the C-terminal carboxylic group are indicated as 'N+' and 'C-' following PROPKA nomenclature. A dotted orange line is also shown at correlation = 0.5.

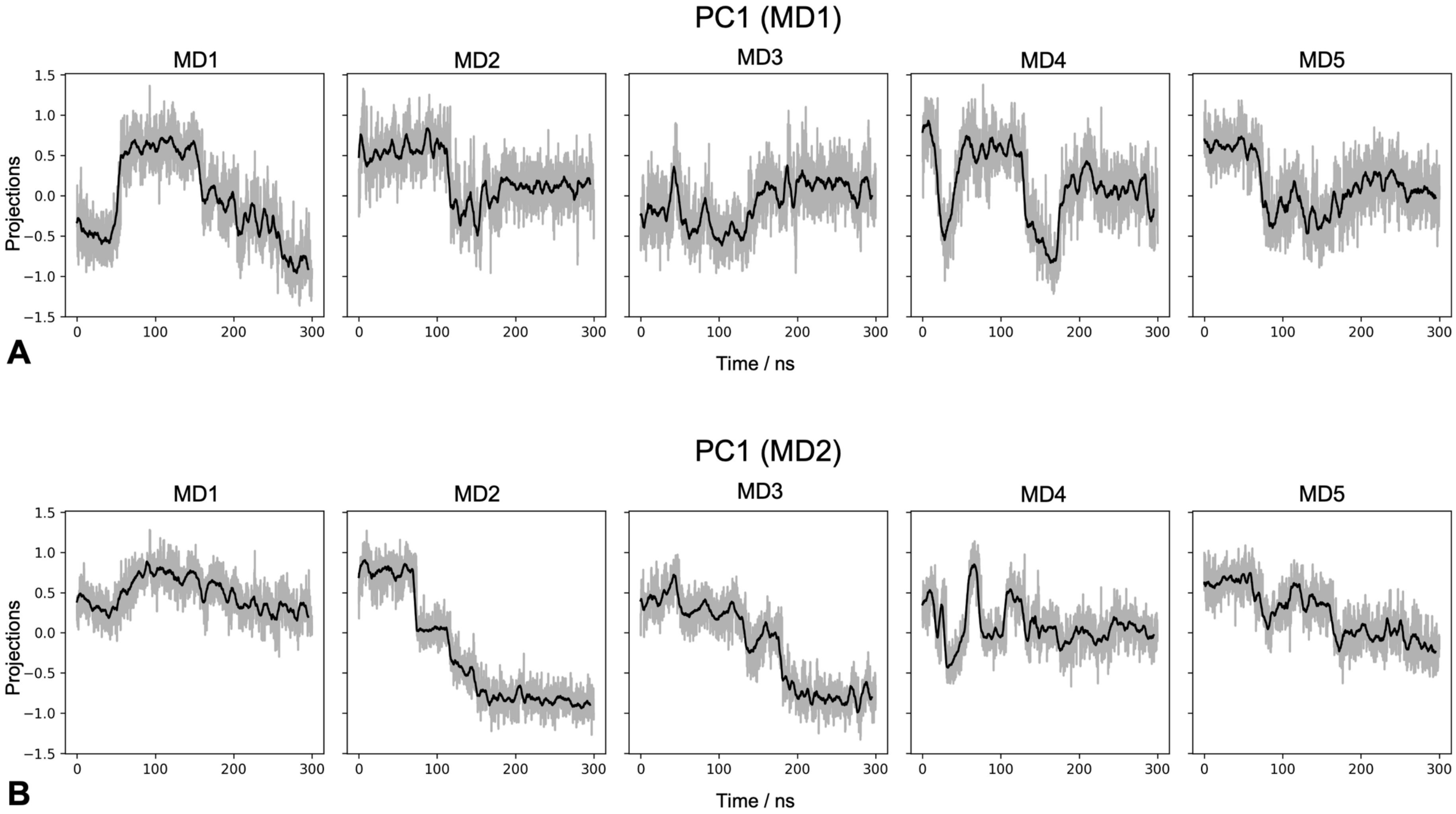


**Figure S4.** Time evolution of the projection of each trajectory on PC1 from MD1 (A) and PC1 from MD2 (B). The running average (5-ns window, black) is shown together with the instantaneous values (grey shading).

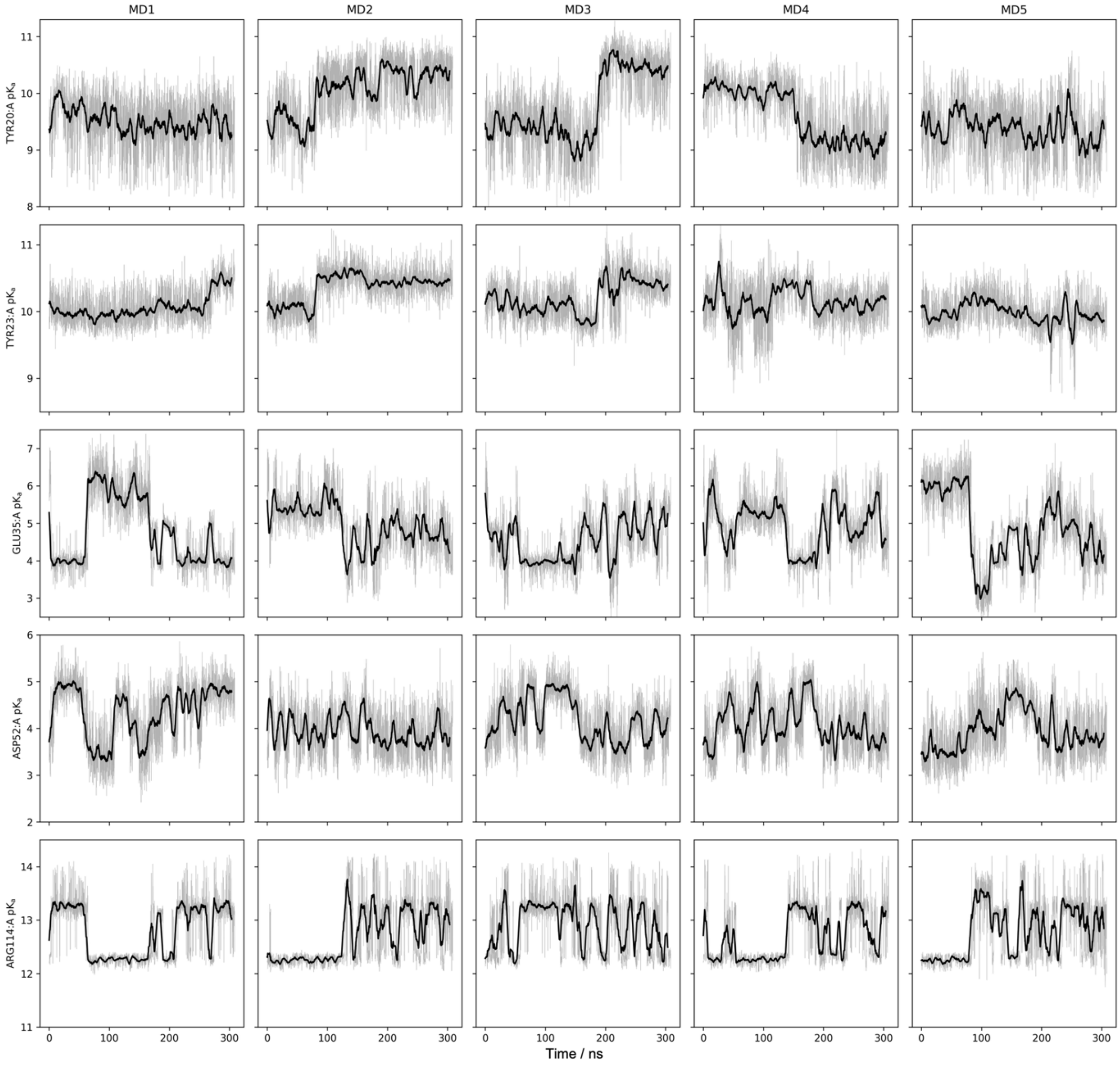


**Figure S5.** Time evolution of p$K_a$ values predicted for Tyr20, Tyr23, Glu35, Asp52 and Arg114 in replicas MD1-5. The running average (5-ns window, dark shade) is shown together with the instantaneous values (light shade).

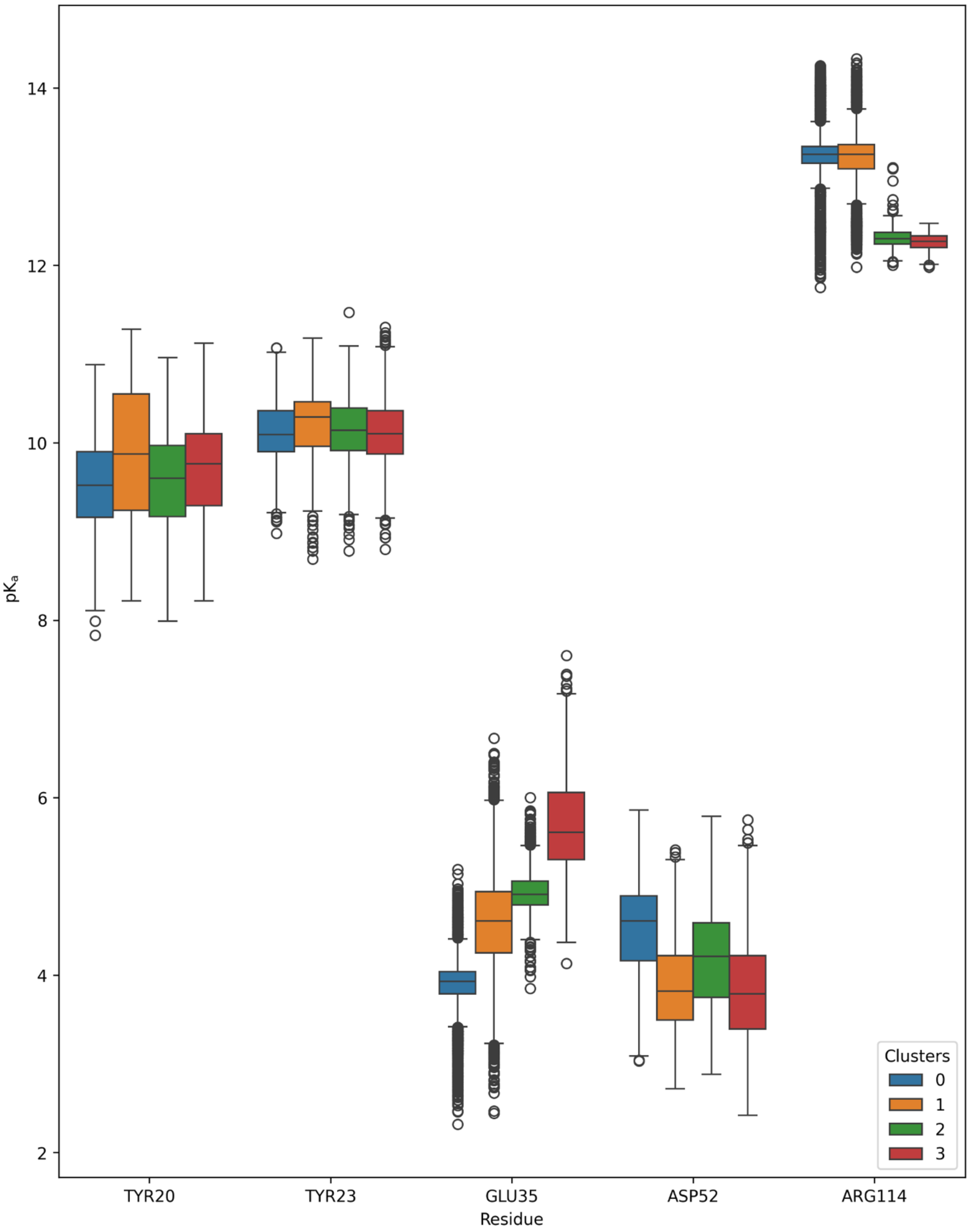


**Figure S6.** Boxplots of p$K_a$ values for Tyr20, Tyr23, Glu35, Asp52 and Arg114 in each of the four K-Medoids clusters.

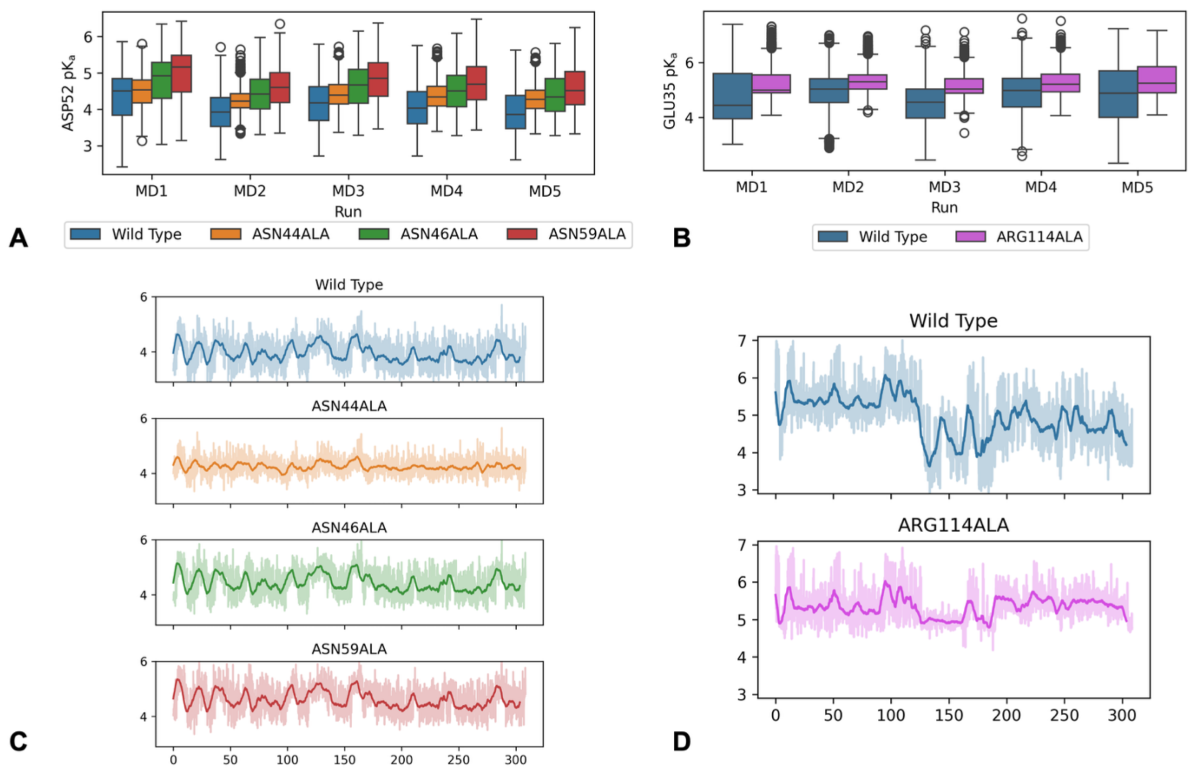


**Figure S7.** (A, B) Boxplots of p$K_a$ values predicted for (A) ASP52 and (B) GLU35 after different pseudomutations for each trajectory. (C, D) Time evolution of p$K_a$ values predicted for (C) ASP52 and (D) GLU35 after different pseudomutations during the MD2 trajectory. The running average (5-ns window, dark shade) is shown together with the instantaneous values (light shade).

REFERENCES


(1) Abraham, M. J.; Murtola, T.; Schulz, R.; Páll, S.; Smith, J. C.; Hess, B.; Lindahl, E. GROMACS: High Performance Molecular Simulations through Multi-Level Parallelism from Laptops to Supercomputers. *SoftwareX* **2015**, *1–2*, 19–25. https://doi.org/10.1016/j.softx.2015.06.001.

(2) Lindahl; Abraham; Hess; Spoel, V. D. GROMACS 2020.3 Manual. **2020**. https://doi.org/10.5281/ZENODO.3923644.

(3) Lindorff-Larsen, K.; Maragakis, P.; Piana, S.; Eastwood, M. P.; Dror, R. O.; Shaw, D. E. Systematic Validation of Protein Force Fields against Experimental Data. *PLoS ONE* **2012**, *7* (2), e32131. https://doi.org/10.1371/journal.pone.0032131.

(4) Artymiuk, P. J.; Blake, C. C. F.; Rice, D. W.; Wilson, K. S. The Structures of the Monoclinic and Orthorhombic Forms of Hen Egg-White Lysozyme at 6 Å Resolution. *Acta Crystallogr B Struct Crystallogr Cryst Chem* **1982**, *38* (3), 778–783. https://doi.org/10.1107/S0567740882004075.

(5) Olsson, M. H. M.; Søndergaard, C. R.; Rostkowski, M.; Jensen, J. H. PROPKA3: Consistent Treatment of Internal and Surface Residues in Empirical p $K_a$ Predictions. *J. Chem. Theory Comput.* **2011**, *7* (2), 525–537. https://doi.org/10.1021/ct100578z.

(6) Jorgensen, W. L.; Chandrasekhar, J.; Madura, J. D.; Impey, R. W.; Klein, M. L. Comparison of Simple Potential Functions for Simulating Liquid Water. *J. Chem. Phys.* **1983**, *79* (2), 926–935. https://doi.org/10.1063/1.445869.

(7) Hess, B.; Bekker, H.; Berendsen, H. J. C. LINCS: A Linear Constraint Solver for Molecular Simulations. *J. Comp. Chem.* **1997**, *18* (12), 1463–1472.

(8) Miyamoto, S.; Kollman P. A. Settle: An Analytical Version of the SHAKE and RATTLE Algorithm for Rigid Water Models. *J. Comp. Chem.* **1992**, *13* (8), 952--962.

(9) Berendsen, H. J. C.; Postma, J. P. M.; Gunsteren, W. F.; Di Nola, A.; Haak, J. R. Molecular Dynamics with Coupling to an External Bath. *J. Chem. Phys.* **1984**, *81* (9), 3684--3690.

(10) Bussi, G.; Donadio, D.; Parrinello, M. Canonical Sampling through Velocity Rescaling. *J. Chem. Phys.* **2007**, *126* (1), 014101. https://doi.org/10.1063/1.2408420.

(11) Parrinello, M.; Rahman, A. Polymorphic Transitions in Single Crystals: A New Molecular Dynamics Method. *J. Appl. Phys.* **1981**, *52* (9), 7182–7190.

(12) Bartik, K.; Redfield, C.; Dobson, C. M. Measurement of the Individual pKa Values of Acidic Residues of Hen and Turkey Lysozymes by Two-Dimensional 1H NMR. *Biophysical Journal* **1994**, *66* (4), 1180–1184. https://doi.org/10.1016/S0006-3495(94)80900-2.

(13) Webb, H.; Tynan-Connolly, B. M.; Lee, G. M.; Farrell, D.; O'Meara, F.; Søndergaard, C. R.; Teilum, K.; Hewage, C.; McIntosh, L. P.; Nielsen, J. E. Remeasuring HEWL p$K_a$ Values by NMR Spectroscopy: Methods, Analysis, Accuracy, and Implications for Theoretical p$K_a$ Calculations. *Proteins* **2011**, *79* (3), 685–702. https://doi.org/10.1002/prot.22886.
